# Identifying common genes, proteins, and pathways from human miRNA and gene blood profiles in multiple sclerosis patients

**DOI:** 10.1101/2022.11.29.518394

**Authors:** Souvik Chakraborty, Tarasankar Maiti, Sushmita Bhowmick, Soumili Sarkar

## Abstract

The molecular pathway associated with Multiple sclerosis (MS) is complex and symptomatic treatments are only available right now. Early diagnosis of MS creates a window for healthcare providers to manage the disease more efficiently. Blood-based biomarker study has been done in the past to identify the upregulated and downregulated genes but in this present study, a novel approach has been taken for identifying genes associated with the disease. In this present study, hub genes are identified and the top ten hub genes were used to identify drugs associated with them. Upregulated genes were identified using the dataset GSE21942 (which contains information related to genes identified in the blood of multiple sclerosis patients) and datasets GSE17846 and GSE61741(which contains information related to microRNAs taken from multiple sclerosis patients). Genes associated with microRNAs were identified using miRWalk. Common genes from both miRWalk and the dataset GSE21942 were identified and were subjected to STRINGdb for the creation of a protein-protein interaction network and this network was then imported to Cytoscape for identifying the top ten hub genes. The top ten hub genes were subjected to EnrichR for enrichment analysis of genes. In our study, it was found that CTNNB1 is the gene with the highest degree (116).

## Introduction

Multiple sclerosis (MS) is an autoimmune ailment associated with demyelination followed by neurodegeneration (Mammana et al., 2018). The frequency of MS on a worldwide scale is 30.1 cases per 100,000 population in 2016 (Dilokthornsakul et al., 2016). The symptoms of MS start at an early age and are more frequently seen in women than men (Wallin et al., 2019). MS is a progressive disease that culminates in the loss of myelin sheath leading to the symptoms such as ataxia, visual complications, fatigue, impaired coordination, and sensation of numbness (Ghasemi et al., 2017). The advancement in the field of magnetic resonance imaging techniques has significantly led to the early detection of the disease (Miller, 1998). In the present study, two datasets contain the microRNA expression profile from the peripheral blood of the patients and normal control subjects. The differentially expressed microRNAs (DEMs) were then subjected to miRWalk, and the genes associated with the DEMs are identified (Dweep et al., 2014). The third dataset of our study (GSE21942) contains information on the genes associated with MS in peripheral blood samples of control and patients. Differentially expressed genes (DEGs) were identified and the common DEGs from the dataset GSE21942 and the DEGs were obtained from miRWalk. The DEGs thus obtained were used for the creation of a protein-protein interaction network (PPIN). The PPIN was then used for the identification of hub genes followed by enrichment analysis of identified hub genes.

## Materials and Methods

### Dataset selection

The microarray gene expression data used in this study were obtained from the GEO (Barrett et al., 2012). The keywords used for the database search are “Multiple sclerosis”, “microRNAs”, “gene expression”, and “expression profiling by array”. Three datasets were chosen for our study. The datasets GSE17846 and GSE61741 used peripheral blood samples for non-coding RNA in MS patients and normal controls (Keller et al., 2009, 2014). We have selected 21 control samples and 20 diseased samples, 23 control samples, and 23 diseased samples respectively. The GSE21942 dataset was used for obtaining gene expression data from the peripheral blood of MS patients (Kemppinen et al., 2011). The dataset used Affymetrix Gene Chip Human Genome U133 Plus 2.0 Array and we have selected 10 normal and 10 MS patients for our study.

### Identification of DEMs and DEGs

All the datasets are analyzed using GEO2R and after analysis, DEMs and DEGs were identified by applying a p-value of 0.05 and the fold change (Log FC) value greater than or equal to 0.5. Common DEMs (in the case of datasets GSE17846 and GSE61741) are then identified using the FunRich tool by creating a Venn diagram (Pathan et al., 2015). After the identification of common DEMs, the DEMs were subjected to miRWalk for the identification of genes associated with the DEMs. The genes associated with DEMs were then subjected to FunRich together with common DEGs associated with the dataset GSE21942 and a Venn diagram was created with the common genes from the dataset and the genes associated with the DEMs.

### Identification of hub genes from PPIN

The common DEGs are subjected to the Search Tool for retrieval of genes (STRING) database and the PPIN was created using medium confidence (Szklarczyk et al., 2019). The PPIN was subjected to Cytoscape and the cytoHubba plugin was used for the identification of hub genes (Chin et al., 2014; Shannon et al., 2003).

### Enrichment analysis of hub genes

The top ten hub genes were subjected to EnrichR webserver for the enrichment analysis of genes (Kuleshov et al., 2016). Appyter notebook was used for the development of bar graphs of significant KEGG pathways associated with the

## Results and discussion

### Identification of common DEGs

In the datasets, GSE17846 and GSE61741, 85 DEMs were identified (Fig 1)

**Fig 1.**
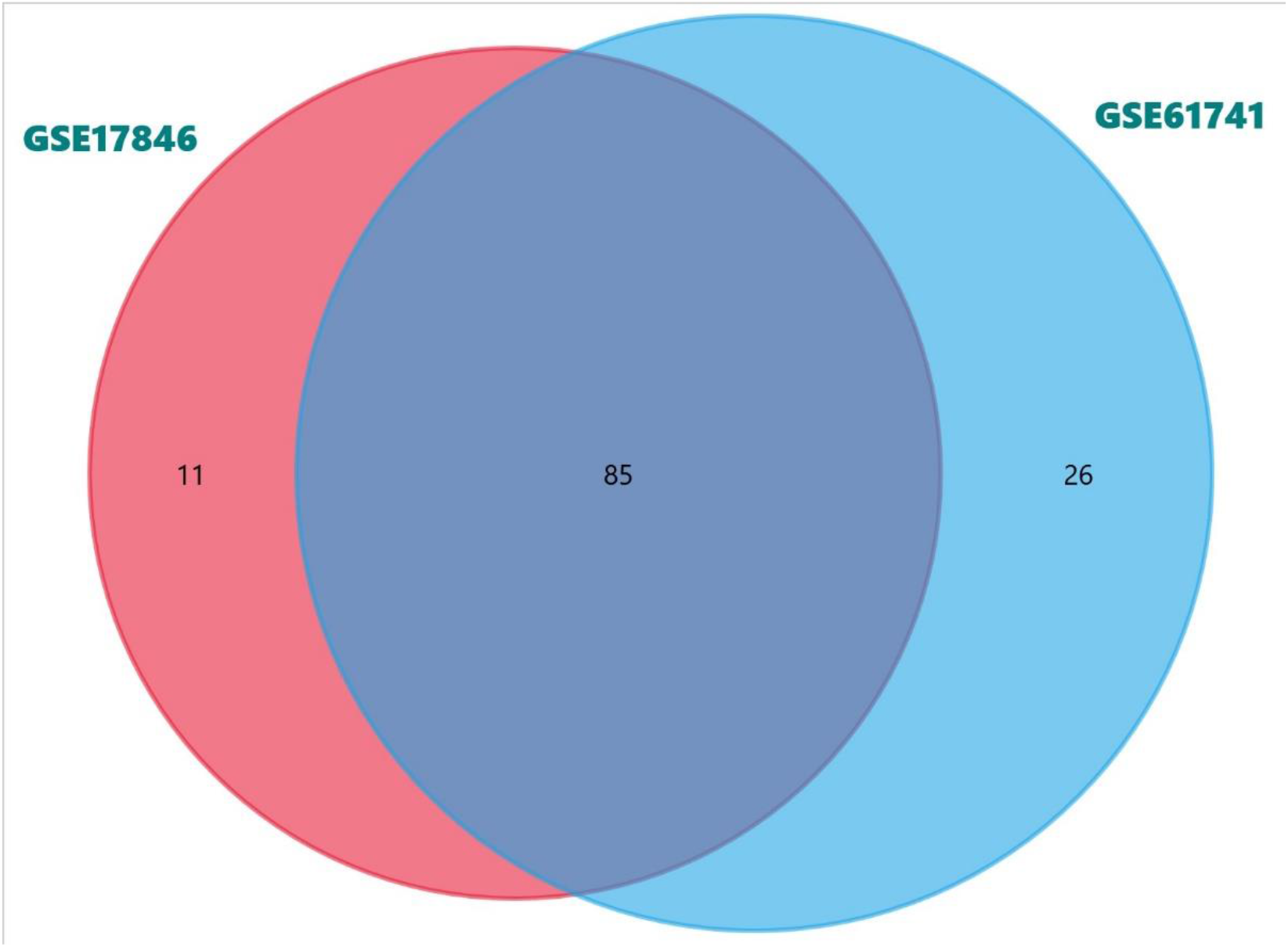
Venn diagram showing DEMs between the datasets GSE17846 and GSE61741.

The 85 DEMs thus obtained are subjected to miRWalk and 15,532 genes are found to be associated. The DEGs between the 15,532 genes and GSE21942 datasets were found to be 915 (Fig 2).

**Fig 2.**
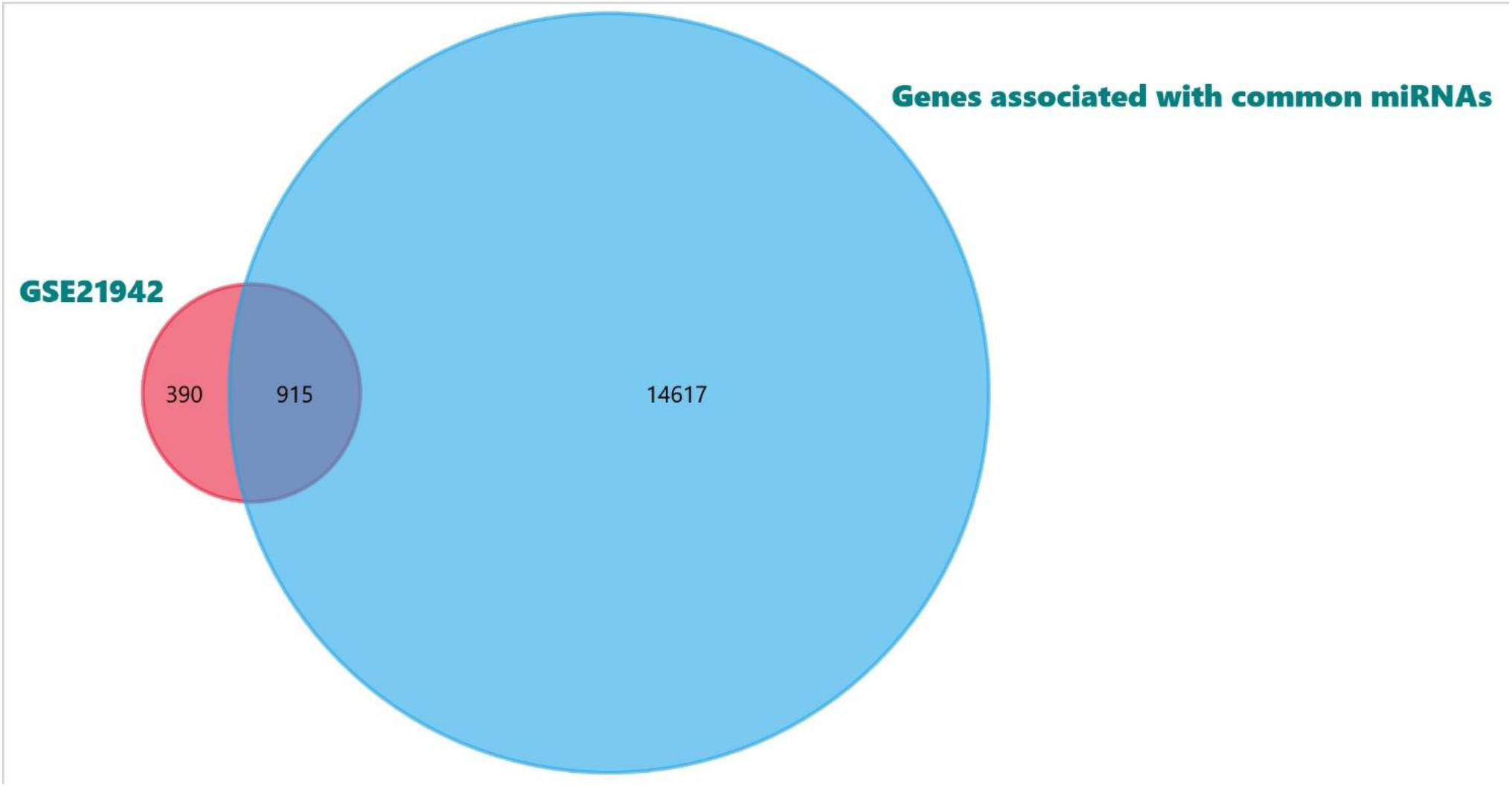
Venn diagram showing the DEGs between the GSE21942 and the genes associated with DEMs obtained from miRWalk.

### Creation of PPIN

These 915 genes were subjected to the STRING database and a PPIN was developed. The PPIN has 914 nodes and 4264 edges with an average node degree of 9.33. The interaction score was set to medium confidence. The entire PPIN is represented in Fig 3.

**Fig 3.**
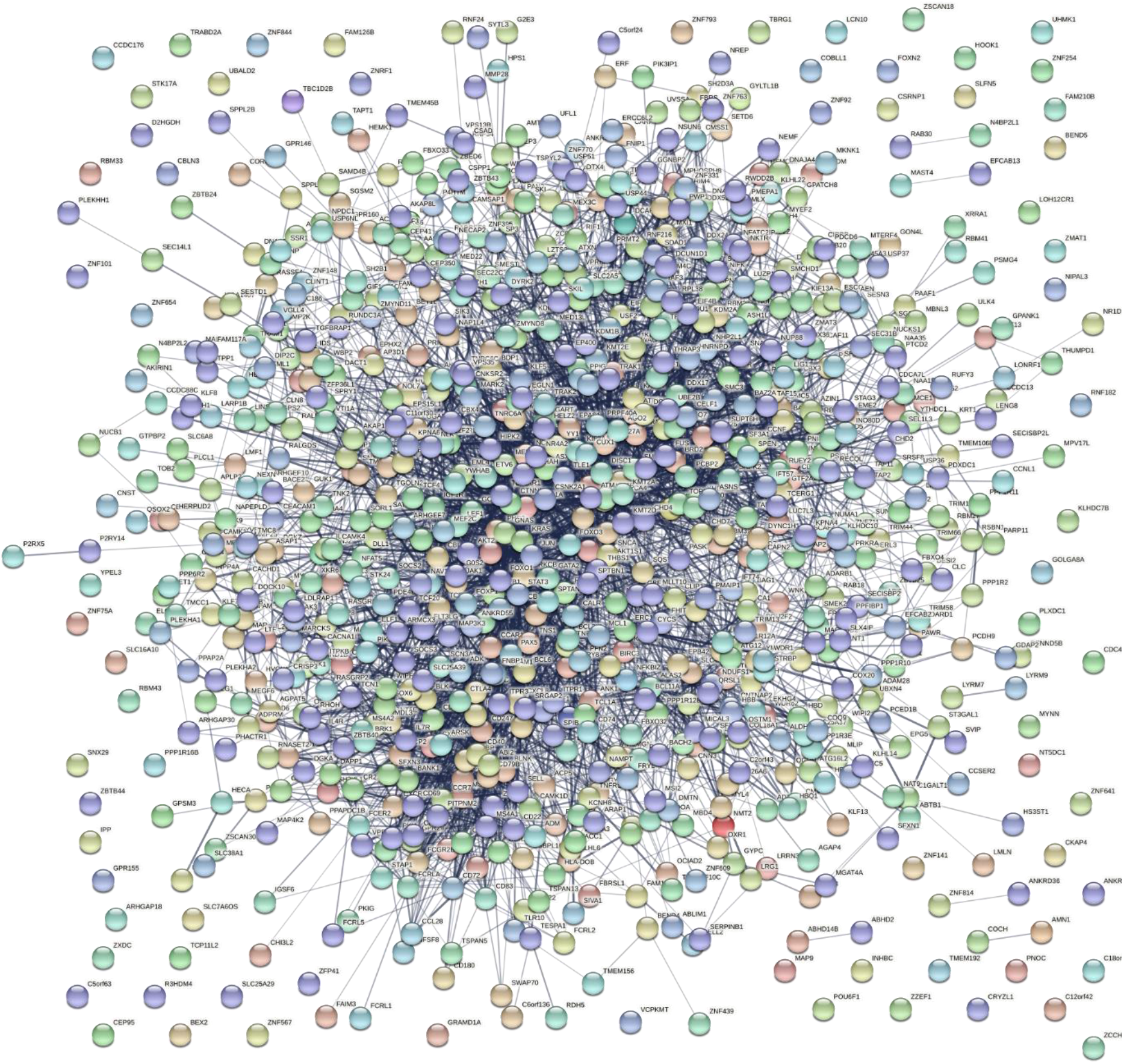
STRING database image showing PPIN of 915 DEGs. In this PPIN the interaction score was set at medium confidence of 0.400.

The above PPIN was imported to Cytoscape software and the top ten hub genes were identified using the CytoHubba plug-in (Fig 4). The gene CTNNB1 had the highest degree of 116 and BCL6 with the lowest degree of 53. The complete list of all the genes with the degree of top ten hub genes is presented in Table 1. The CTNNB1 gene encodes beta-catenin which is important for Wnt/ beta-catenin pathway. Activation of the wnt/beta-catenin pathway is associated with the loss of remyelination (Gaesser & Fyffe-Maricich, 2016).

**Fig 4.**
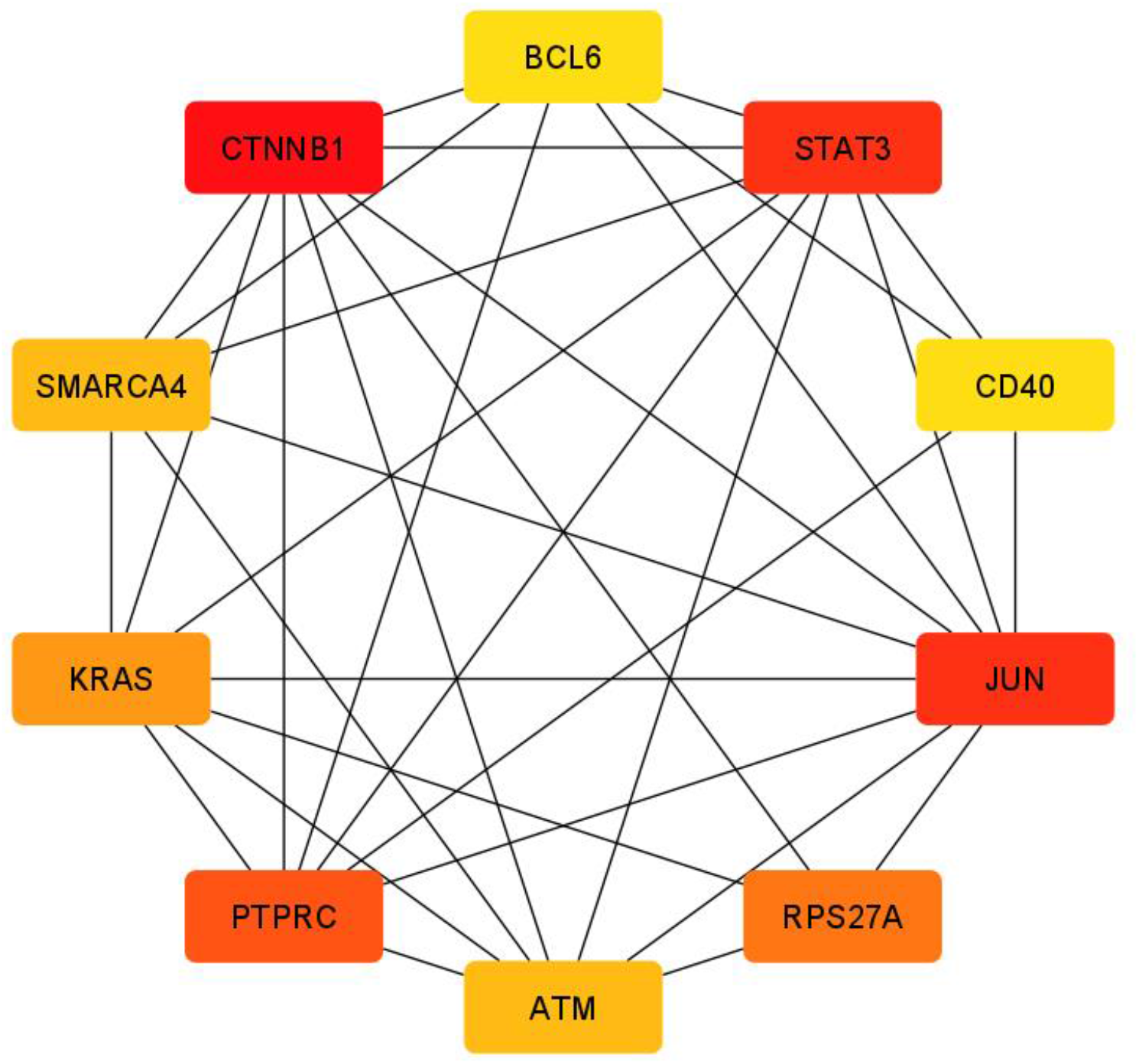
Network of top ten hub genes. The red colour in this represents the maximum degree of centrality, orange represents the medium degree of centrality and yellow represents the lowest degree of centrality.

**Table 1:** Top ten Hub proteins in the PPI network obtained from the DEGs of the datasets for HD and MS with their degree of centrality.

| Gene | Degree |
| --- | --- |
| CTNNB1 | 116 |
| JUN | 90 |
| STAT3 | 90 |
| PTPRC | 74 |
| RPS27A | 71 |
| KRAS | 69 |
| ATM | 62 |
| SMARCA4 | 62 |
| CD40 | 53 |
| BCL6 | 53 |

Activation of Jun transcription factor has been associated with MS (Bonetti et al., 1999). STAT 3 is essential for the regulaton of balance between T effector and T regulatory cells which is hampered in MS. Inhibition of STAT3 by inhibitor have been found to be beneficial for patients with relapsing remitting type MS which shows that STAT3 can be used as a potential therapeutic target in the future for MS (Aqel et al., 2021). PTPRC a gene that encodes a protein called CD45 has not been connected to MS yet but more extensive studies are now required to confirm the above statement (Miterski et al., 2002). In a study conducted by Chen et al. it was found out that RPS27A is one of the hub genes in a network of genes which are upregulated (Chen et al., 2021). Ras proteins are major player in the development of human cancers but it was also found that these proteins play a role in the development of MS. Inhibition of Ras protein directly has not been a successful strategy but recently repurposing of cancer drugs for selectively inhibiting the KRAS may provide an answer to the long standing question but more research is required on this matter (Sun et al., 2012).

Impaired activation of ATM leads to impaired activation of p53 protein thus leading to apoptotic pathway activation in MS patients (Deng et al., 2005). Genetic mutations in SMARC genes has been associated with high prevalence of MS (Valori et al., 2021). Lower expression of CD40 has been associated with high risk of MS (Field et al., 2015). Bcl6 plays an important role in controlling IL6 secretion from macrophage and IL6 has been significantly associated with demyelination in CNS and thus can play major role in the development of MS but more studies are required on this (Schneider et al., 2013).

### Identification of KEGG pathways enriched in top ten hub genes

The top ten hub genes were subjected to Enrichr and significant KEGG pathways, common GO biological processes, and GO molecular function.

### Common enriched KEGG pathways

The Kaposi sarcoma-associated herpes virus infection pathways were enriched in the gene set uploaded. A study showed that the human herpes virus 8 genomes were detected in 10 out of 54 individuals with MS and 4 out of 130 control samples when nested PCR was performed (Marashi et al., 2018). Forkhead box O1 in short FoxO signaling pathway was also enriched in the geneset. Inhibition of Fox O1 results in a decrease in the autoimmune response of T cells (Kraus et al., 2021). In a study, it was found that participants with relapsing-remitting type MS also was affected by dyslipidemia, and thus it goes with our finding that pathways for atherosclerosis are enriched in our gene set (Etemadifar et al., 2022). There was no evidence of human T cell leukemia virus 1 infection with multiple sclerosis. Several cancers like breast cancer, gastrointestinal cancers, and central nervous system cancers have been associated with a higher risk of MS and go per the findings (Catalá-López et al., 2014; Nielsen et al., 2006). The mitophagy and colorectal cancer pathways are associated with MS and in a study, it was found that the survival of patients with colorectal cancer was low compared to non-MS patients (Giorgi et al., 2021; Marrie et al., 2021). Programmed cell death ligand 1 or PDL1 has been associated with MS (Mi et al., 2021). According to a recent study, the role of advanced glycation end products (AGE) and their receptors (RAGE) has been studied in MS patients but the study could not conclude that there is a direct role of AGE and RAGE in MS (Damasiewicz-Bodzek et al., 2021). MS is an autoimmune disease and studies state that T cell receptor signaling has been directly linked to MS pathogenesis (van Langelaar et al., 2020). The KEGG pathways associated with the top ten hub genes have been shown in figure 5.

**Fig 5.**
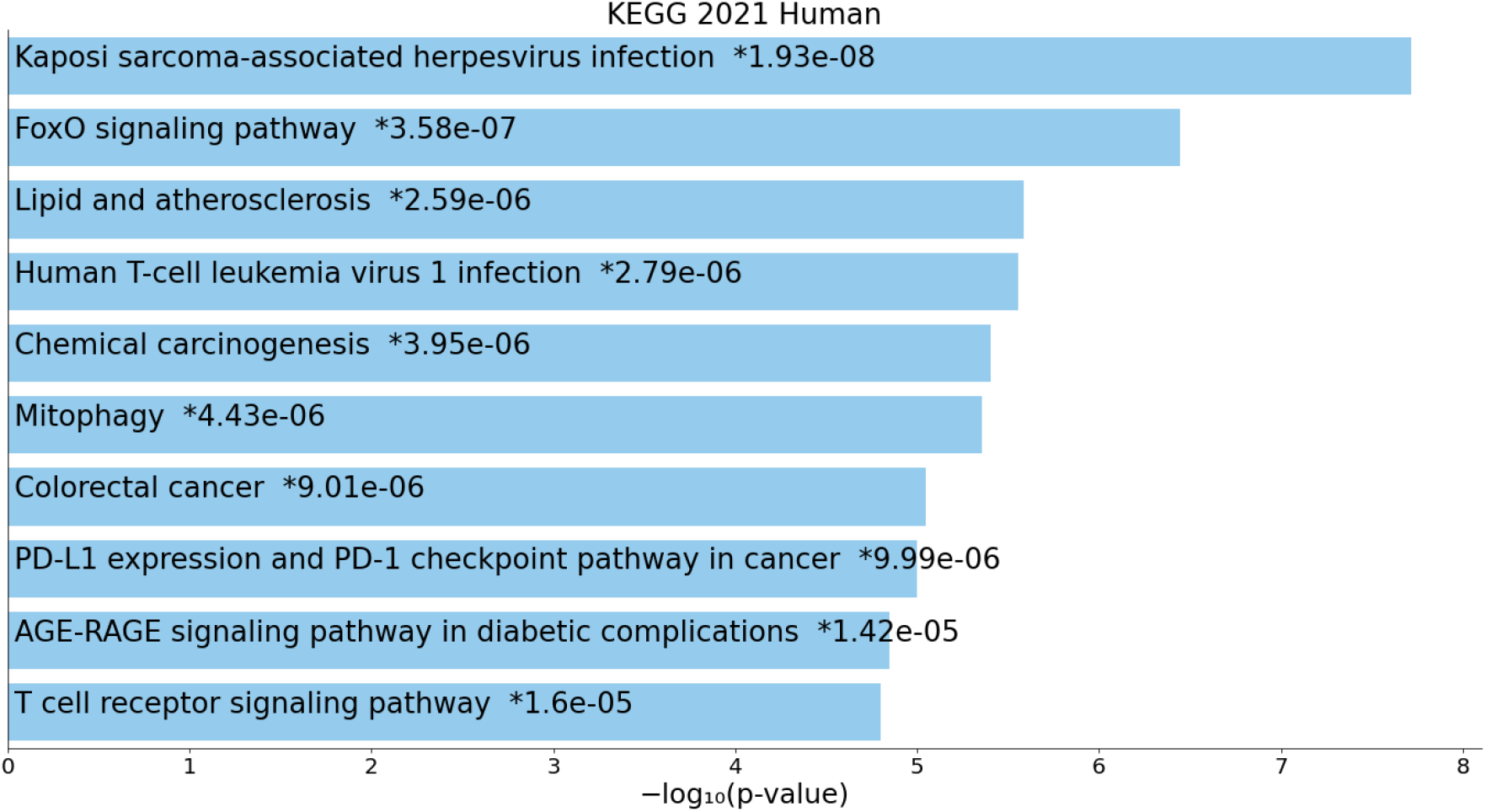
Bar graphs showing the top ten enriched KEGG pathways associated with the top ten hub genes. The x-axis represents the -log_10_(p-value) and the y-axis represents the KEGG pathways associated with the top ten hub genes.

### Common enriched biological processes

The biological processes that are significantly enriched are transcription by RNA polymerase II (GO:0045944) with a p-value of 4.14e-08 and DNA binding transcription factor activity (GO:0051091) with a p-value of 6.48e-08. The graphical representation of enriched biological processes is presented in figure 6.

**Fig 6.**
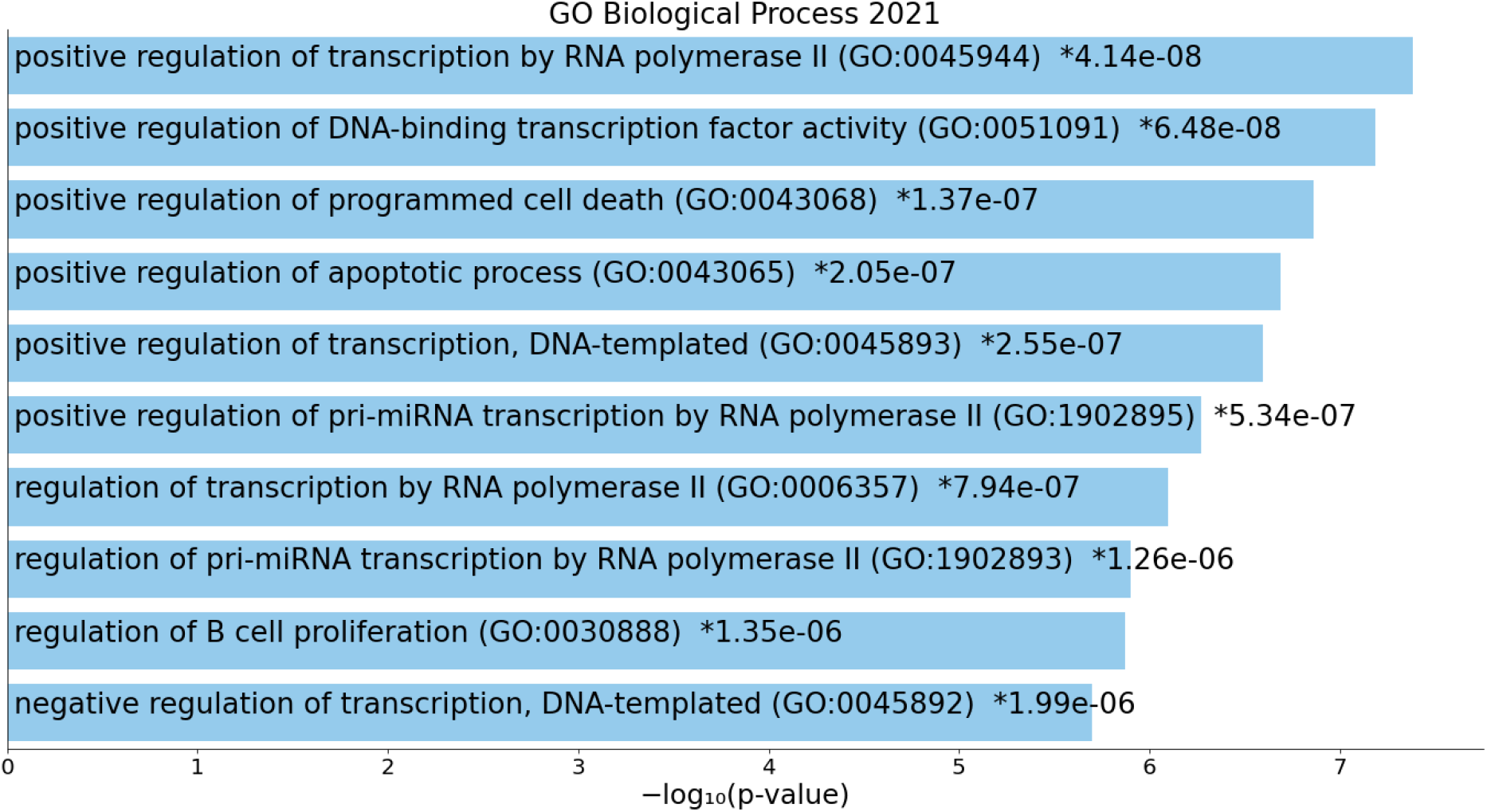
Bar graphs showing the top ten enriched biological processes associated with the top ten hub genes. The x-axis represents the -log_10_(p-value) and the y-axis represents the biological processes associated with the top ten hub genes.

### Common enriched molecular functions

The molecular functions that are significantly enriched are RNA polymerase II specific DNA binding transcription factor binding (GO:0061629) with a p-value of 1.58e-06. The graphical representation of enriched molecular functions is presented in figure 7.

**Fig 7.**
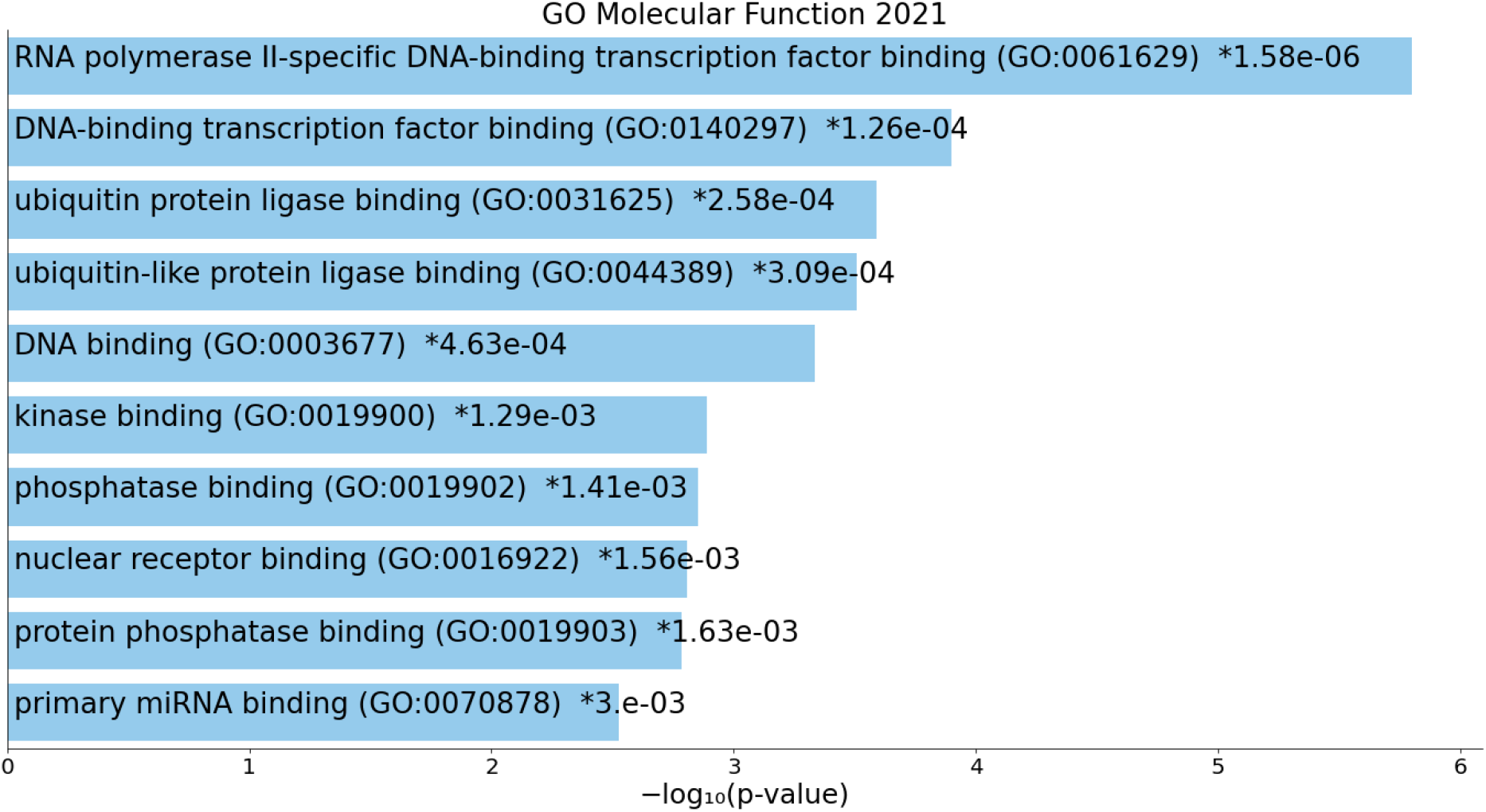
Bar graphs showing the top ten enriched molecular functions associated with the top ten hub genes. The x-axis represents the -log_10_(p-value) and the y-axis represents the molecular functions associated with the top ten hub genes.

Like every study our study also has some drawbacks such a s in our study we have not considered the gender of the subject, we also have not discussed the primary type o MS i.e whether we wanted to target relapsing-remitting type MS or others, and finally, more datasets could have been included in the study.

## Conclusion

In this present study, we have tried to find out a possible biomarker for MS using microarray datasets. We have identified in this study that CTNNB1 or beta-catenin protein is having the highest degree and they have also been associated with MS in earlier studies thus more studies are required to ascertain whether this gene product can become a biomarker for MS or not. In our study, we also found that the most enriched KEGG pathway is Kaposi sarcoma-associated herpes virus infection thus leading to the conclusion that the common proteins associated with MS and the aforesaid infection should be carefully examined for finding therapeutic agents for the cure of MS in the rare future.

## Supporting information

https://drive.google.com/file/d/1DFTwZN4IpuRyomevhA3yzAF2hPsx__fl/view?usp=share_link

## References

Aqel, S. I., Yang, X., Kraus, E. E., Song, J., Farinas, M. F., Zhao, E. Y., Pei, W., Lovett-Racke, A. E., Racke, M. K., Li, C., & Yang, Y. (2021). A STAT3 inhibitor ameliorates CNS autoimmunity by restoring Teff:Treg Balance. JCI Insight. https://doi.org/10.1172/jci.insight.142376

Barrett, T., Wilhite, S. E., Ledoux, P., Evangelista, C., Kim, I. F., Tomashevsky, M., Marshall, K. A., Phillippy, K. H., Sherman, P. M., Holko, M., Yefanov, A., Lee, H., Zhang, N., Robertson, C. L., Serova, N., Davis, S., & Soboleva, A. (2012). NCBI GEO: Archive for functional genomics data sets—update. Nucleic Acids Research, 41(D1), D991–D995. https://doi.org/10.1093/nar/gks1193

Bonetti, B., Stegagno, C., Cannella, B., Rizzuto, N., Moretto, G., & Raine, C. S. (1999). Activation of NF-κB and c-jun Transcription Factors in Multiple Sclerosis Lesions. The American Journal of Pathology, 155(5), 1433–1438. https://doi.org/10.1016/S0002-9440(10)65456-9

Catalá-López, F., Suárez-Pinilla, M., Suárez-Pinilla, P., Valderas, J. M., Gómez-Beneyto, M., Martinez, S., Balanzá-Martínez, V., Climent, J., Valencia, A., McGrath, J., Crespo-Facorro, B., Sanchez-Moreno, J., Vieta, E., & Tabarés-Seisdedos, R. (2014). Inverse and Direct Cancer Comorbidity in People with Central Nervous System Disorders: A Meta-Analysis of Cancer Incidence in 577,013 Participants of 50 Observational Studies. Psychotherapy and Psychosomatics, 83(2), 89–105. https://doi.org/10.1159/000356498

Chen, X., Hou, H., Qiao, H., Fan, H., Zhao, T., & Dong, M. (2021). Identification of blood-derived candidate gene markers and a new 7-gene diagnostic model for multiple sclerosis. Biological Research, 54(1), 12. https://doi.org/10.1186/s40659-021-00334-6

Chin, C.-H., Chen, S.-H., Wu, H.-H., Ho, C.-W., Ko, M.-T., & Lin, C.-Y. (2014). cytoHubba: Identifying hub objects and sub-networks from complex interactome. BMC Systems Biology, 8(S4), S11. https://doi.org/10.1186/1752-0509-8-S4-S11

Damasiewicz-Bodzek, A., Łabuz-Roszak, B., Kumaszka, B., Tadeusiak, B., & Tyrpień-Golder, K. (2021). The Assessment of Serum Concentrations of AGEs and Their Soluble Receptor (sRAGE) in Multiple Sclerosis Patients. Brain Sciences, 11(8), 1021. https://doi.org/10.3390/brainsci11081021

Deng, X., Ljunggren-Rose, A., Maas, K., & Sriram, S. (2005). Defective ATM-p53-mediated apoptotic pathway in multiple sclerosis. Annals of Neurology, 58(4), 577–584. https://doi.org/10.1002/ana.20600

Dilokthornsakul, P., Valuck, R. J., Nair, K. V., Corboy, J. R., Allen, R. R., & Campbell, J. D. (2016). Multiple sclerosis prevalence in the United States commercially insured population. Neurology, 86(11), 1014–1021. https://doi.org/10.1212/WNL.0000000000002469

Dweep, H., Gretz, N., & Sticht, C. (2014). MiRWalk Database for miRNA–Target Interactions. In M. L. Alvarez & M. Nourbakhsh (Eds.), RNA Mapping (Vol. 1182, pp. 289–305). Springer New York. https://doi.org/10.1007/978-1-4939-1062-5_25

Etemadifar, M., Salari, M., Esnaashari, A., Ghazanfaripoor, F., Sayahi, F., Akhavan Sigari, A., & Sedaghat, N. (2022). Atherosclerosis and multiple sclerosis: An overview on the prevalence of risk factors. Multiple Sclerosis and Related Disorders, 58, 103488. https://doi.org/10.1016/j.msard.2022.103488

Field, J., Shahijanian, F., Schibeci, S., Australia and New Zealand MS Genetics Consortium (ANZgene), Johnson, L., Gresle, M., Laverick, L., Parnell, G., Stewart, G., McKay, F., Kilpatrick, T., Butzkueven, H., & Booth, D. (2015). The MS Risk Allele of CD40 Is Associated with Reduced Cell-Membrane Bound Expression in Antigen Presenting Cells: Implications for Gene Function. PLOS ONE, 10(6), e0127080. https://doi.org/10.1371/journal.pone.0127080

Gaesser, J. M., & Fyffe-Maricich, S. L. (2016). Intracellular signaling pathway regulation of myelination and remyelination in the CNS. Experimental Neurology, 283, 501–511. https://doi.org/10.1016/j.expneurol.2016.03.008

Ghasemi, N., Razavi, S., & Nikzad, E. (2017). Multiple Sclerosis: Pathogenesis, Symptoms, Diagnoses and Cell-Based Therapy. Cell J (Yakhteh), 19(1). https://doi.org/10.22074/cellj.2016.4867

Giorgi, C., Bouhamida, E., Danese, A., Previati, M., Pinton, P., & Patergnani, S. (2021). Relevance of Autophagy and Mitophagy Dynamics and Markers in Neurodegenerative Diseases. Biomedicines, 9(2), 149. https://doi.org/10.3390/biomedicines9020149

Keller, A., Leidinger, P., Lange, J., Borries, A., Schroers, H., Scheffler, M., Lenhof, H.-P., Ruprecht, K., & Meese, E. (2009). Multiple Sclerosis: MicroRNA Expression Profiles Accurately Differentiate Patients with Relapsing-Remitting Disease from Healthy Controls. PLoS ONE, 4(10), e7440. https://doi.org/10.1371/journal.pone.0007440

Keller, A., Leidinger, P., Vogel, B., Backes, C., ElSharawy, A., Galata, V., Mueller, S. C., Marquart, S., Schrauder, M. G., Strick, R., Bauer, A., Wischhusen, J., Beier, M., Kohlhaas, J., Katus, H. A., Hoheisel, J., Franke, A., Meder, B., & Meese, E. (2014). MiRNAs can be generally associated with human pathologies as exemplified for miR-144*. BMC Medicine, 12(1), 224. https://doi.org/10.1186/s12916-014-0224-0

Kemppinen, A. K., Kaprio, J., Palotie, A., & Saarela, J. (2011). Systematic review of genome-wide expression studies in multiple sclerosis. BMJ Open, 1(1), e000053– e000053. https://doi.org/10.1136/bmjopen-2011-000053

Kraus, E. E., Kakuk-Atkins, L., Farinas, M. F., Jeffers, M., Lovett-Racke, A. E., & Yang, Y. (2021). Regulation of autoreactive CD4 T cells by FoxO1 signaling in CNS autoimmunity. Journal of Neuroimmunology, 359, 577675. https://doi.org/10.1016/j.jneuroim.2021.577675

Kuleshov, M. V., Jones, M. R., Rouillard, A. D., Fernandez, N. F., Duan, Q., Wang, Z., Koplev, S., Jenkins, S. L., Jagodnik, K. M., Lachmann, A., McDermott, M. G., Monteiro, C. D., Gundersen, G. W., & Ma’ayan, A. (2016). Enrichr: A comprehensive gene set enrichment analysis web server 2016 update. Nucleic Acids Research, 44(W1), W90–W97. https://doi.org/10.1093/nar/gkw377

Mammana, S., Fagone, P., Cavalli, E., Basile, M., Petralia, M., Nicoletti, F., Bramanti, P., & Mazzon, E. (2018). The Role of Macrophages in Neuroinflammatory and Neurodegenerative Pathways of Alzheimer’s Disease, Amyotrophic Lateral Sclerosis, and Multiple Sclerosis: Pathogenetic Cellular Effectors and Potential Therapeutic Targets. International Journal of Molecular Sciences, 19(3), 831. https://doi.org/10.3390/ijms19030831

Marashi, S. M., Mostafa, A., Shoja, Z., Nejati, A., Shahmahmoodi, S., Mollaei-Kandelous, Y., Sahraian, M. A., & Jalilvand, S. (2018). Human herpesvirus 8 DNA detection and variant analysis in patients with multiple sclerosis. VirusDisease, 29(4), 540–543. https://doi.org/10.1007/s13337-018-0481-1

Marrie, R. A., Maxwell, C., Mahar, A., Ekuma, O., McClintock, C., Seitz, D., & Groome, P. (2021). Colorectal Cancer Survival in Multiple Sclerosis: A Matched Cohort Study. Neurology, 97(14), e1447–e1456. https://doi.org/10.1212/WNL.0000000000012634

Mi, Y., Han, J., Zhu, J., & Jin, T. (2021). Role of the PD-1/PD-L1 Signaling in Multiple Sclerosis and Experimental Autoimmune Encephalomyelitis: Recent Insights and Future Directions. Molecular Neurobiology, 58(12), 6249–6271. https://doi.org/10.1007/s12035-021-02495-7

Miller, D. (1998). The role of magnetic resonance techniques in understanding and managing multiple sclerosis. Brain, 121(1), 3–24. https://doi.org/10.1093/brain/121.1.3

Miterski, B., Sindern, E., Haupts, M., Schimrigk, S., & Epplen, J. T. (2002). PTPRC(CD45) is not associated with multiple sclerosis in a large cohort of German patients. BMC Medical Genetics, 3(1), 3. https://doi.org/10.1186/1471-2350-3-3

Nielsen, N. M., Rostgaard, K., Rasmussen, S., Koch-Henriksen, N., Storm, H. H., Melbye, M., & Hjalgrim, H. (2006). Cancer risk among patients with multiple sclerosis: A population-based register study. International Journal of Cancer, 118(4), 979–984. https://doi.org/10.1002/ijc.21437

Pathan, M., Keerthikumar, S., Ang, C.-S., Gangoda, L., Quek, C. Y. J., Williamson, N. A., Mouradov, D., Sieber, O. M., Simpson, R. J., Salim, A., Bacic, A., Hill, A. F., Stroud, D. A., Ryan, M. T., Agbinya, J. I., Mariadason, J. M., Burgess, A. W., & Mathivanan, S. (2015). FunRich: An open access standalone functional enrichment and interaction network analysis tool. PROTEOMICS, 15(15), 2597–2601. https://doi.org/10.1002/pmic.201400515

Schneider, A., Long, S. A., Cerosaletti, K., Ni, C. T., Samuels, P., Kita, M., & Buckner, J. H. (2013). In Active Relapsing-Remitting Multiple Sclerosis, Effector T Cell Resistance to Adaptive T _regs_ Involves IL-6–Mediated Signaling. Science Translational Medicine, 5(170). https://doi.org/10.1126/scitranslmed.3004970

Shannon, P., Markiel, A., Ozier, O., Baliga, N. S., Wang, J. T., Ramage, D., Amin, N., Schwikowski, B., & Ideker, T. (2003). Cytoscape: A Software Environment for Integrated Models of Biomolecular Interaction Networks. Genome Research, 13(11), 2498–2504. https://doi.org/10.1101/gr.1239303

Sun, Q., Burke, J. P., Phan, J., Burns, M. C., Olejniczak, E. T., Waterson, A. G., Lee, T., Rossanese, O. W., & Fesik, S. W. (2012). Discovery of Small Molecules that Bind to K-Ras and Inhibit Sos-Mediated Activation. Angewandte Chemie International Edition, 51(25), 6140–6143. https://doi.org/10.1002/anie.201201358

Szklarczyk, D., Gable, A. L., Lyon, D., Junge, A., Wyder, S., Huerta-Cepas, J., Simonovic, M., Doncheva, N. T., Morris, J. H., Bork, P., Jensen, L. J., & Mering, C. von. (2019). STRING v11: Protein–protein association networks with increased coverage, supporting functional discovery in genome-wide experimental datasets. Nucleic Acids Research, 47(D1), D607–D613. https://doi.org/10.1093/nar/gky1131

Valori, M., Jansson, L., & Tienari, P. J. (2021). CD8+ cell somatic mutations in multiple sclerosis patients and controls—Enrichment of mutations in STAT3 and other genes implicated in hematological malignancies. PLOS ONE, 16(12), e0261002. https://doi.org/10.1371/journal.pone.0261002

van Langelaar, J., Rijvers, L., Smolders, J., & van Luijn, M. M. (2020). B and T Cells Driving Multiple Sclerosis: Identity, Mechanisms and Potential Triggers. Frontiers in Immunology, 11, 760. https://doi.org/10.3389/fimmu.2020.00760

Wallin, M. T., Culpepper, W. J., Nichols, E., Bhutta, Z. A., Gebrehiwot, T. T., Hay, S. I., Khalil, I. A., Krohn, K. J., Liang, X., Naghavi, M., Mokdad, A. H., Nixon, M. R., Reiner, R. C., Sartorius, B., Smith, M., Topor-Madry, R., Werdecker, A., Vos, T., Feigin, V. L., & Murray, C. J. L. (2019). Global, regional, and national burden of multiple sclerosis 1990–2016: A systematic analysis for the Global Burden of Disease Study 2016. The Lancet Neurology, 18(3), 269–285. https://doi.org/10.1016/S1474-4422(18)30443-5

